# Tail Vein Injections of Recombinant Human Thioredoxin Prevents High Fat-induced Endothelial Dysfunction in Mice

**DOI:** 10.1101/694083

**Authors:** Rob H.P. Hilgers

## Abstract

**Background:** Obesity is a serious risk factor for cardiovascular diseases. A high fat diet results in cellular oxidative stress and endothelial dysfunction in resistance-sized arteries, characterized by reduced nitric oxide (NO) and endothelium-dependent hyperpolarizing (EDH) responses. Thioredoxin-1, a sulfo-oxidoreductase protein that cleaves disulfide bridges between two adjacent cysteine residues in oxidized proteins, has been shown to lower blood pressure and improve endothelium-dependent relaxing responses in aged C57Bl6/J mice.

**Methods and Results:** Young (∼ 3 month-old) male C57Bl6/J mice were fed a high fat diet (42% kcal from fat; obese) or a normal chow (lean) for 3 months. Mice were administered recombinant human thioredoxin-1 (rhTrx; 25 mg/kg) or saline (0.9% NaCl) via tail vein injection at the start, after one month, and after two months. Body weight (BW) was comparable between lean/rhTrx1 and lean/saline at the time of euthanasia (32 ±1 g versus 32 ± 1 g). The high fat regimen resulted in a comparable BW between obese/saline and obese/rhTrx mice (47 ± 1 g versus 45 ± 2 g, respectively). Small (second-order branches) mesenteric arteries (MA2), coronary and femoral arteries were isolated and mounted on the wire-myograph. MA2 and femoral arteries from obese/saline had blunted acetylcholine (10^−9^ – 10^−5^ M)-mediated relaxations compared to lean/saline mice, but not to the NO donor sodium nitroprusside. NO and EDH-mediated relaxing responses were blunted in MA2 from obese/lean mice compared to the three other groups.

**Conclusion:** Tail vein injections with rhTrx prevented endothelial dysfunction in obese mice by improving NO and EDH relaxing responses in MA2.

---

In resistance-sized arteries (< 250 μm in lumen diameter) relaxation is mediated via endothelium-derived factors such as NO and prostacyclin (PGI_2_), but also by a conductive electro-coupled pathway, called endothelium-dependent hyperpolarization (EDH), that is associated with a propagation of endothelial and smooth muscle cell hyperpolarization [1]. Endothelial calcium-activated potassium channels (K_Ca_) initiate and propagate this EDH response [1]. In the endothelium of small mesenteric arteries, small-conductance K_Ca_ (SK_Ca_) and intermediate-conductance K_Ca_ (IK_Ca_) are solely responsible for this EDH [2]. In isometric wire-myography this EDH can be assessed in a contracted artery segment followed by endothelial K_Ca_ opening under conditions where NO and PGI_2_ release is inhibited. Under pathological conditions, including obesity, the contribution of these endothelium-derived mediators is compromised (for a review see [3, 4]). An altered cellular reduction-oxidation (redox) balance shifted towards a more oxidative state is presumed to be the culprit of this endothelial dysfunction [5, 6]. Thiol/disulfide redox changes of specific amino acids, most notably cysteine, modulate the activity of many enzymes [7]. Thioredoxin-1 (Trx) is a 12-kDa cytosolic oxidoreductase capable of reducing disulfide bridges between two adjacent cysteine residues, hereby keeping cysteine groups in its active thiol (reduced) formation [8]. Overexpression of human Trx in mice has been elegantly shown to reduce age-related hypertension [9], to increase endothelium-dependent acetylcholine-mediated relaxations in the mesenteric vascular bed [9, 10] and to preserve endothelial nitric oxide synthase (eNOS) activity via a reductive deglutathionylation process [11]. In addition, the EDH response was enhanced in small mesenteric arteries derived from these Trx transgenic mice compared to their wild-type littermates [10]. These observations suggest that Trx maintains the activity of eNOS and endothelial K_Ca_ channels. In this study it was hypothesized that tail vein injections of recombinant human Trx (rhTrx) would protect against a high fat diet-induced impairment in endothelium-dependent relaxation in resistance-sized arteries in C57Bl6/J mice. In this study, attention was focused on the role of endothelium-derived NO and EDH relaxing responses in mediating ACh- and NS309-induced relaxations in murine arteries derived from lean and obese mice with or without intervention with rhTrx.

## Methods

### Animals and tail vein injections

Male C57Bl6/J mice (10 – 12 weeks) were placed on either a normal chow (lean group) or a high fat diet (obese group). The high fat (42% kcal from fat) diet was purchased from Harlan Laboratories (Teklad Custom Research Diet TD.88137). Mice were divided in four groups: lean/saline, obese/saline, lean/rhTrx, and obese/rhTrx. In the saline groups, mice were injected via the tail vein with saline (100 μL of a 0.9% NaCl sterile solution), and for the rhTrx group with recombinant human Thioredoxin-1 (R&D Systems; 2.5 mg/kg in 100 μL solution in 0.9% NaCl solution). Mice were briefly placed in a holding chamber and anesthetized with isoflurane (1.5% delivered in 100% O_2_). The tail was heated with a light source in order to dilate the tail vein. Tail vein injections were performed with insulin syringes (Exel, 30G). After one and two months the tail vein injections were repeated. After three months mice were euthanized. All procedures were approved by the IACUC at Campbell University and were consistent with the *Guide for the Care and Use of Laboratory Animals* published by the National Institute of Health. All animals were maintained on a standard 12-h light/12-h dark cycle, in a temperature-controlled barrier facility.

### Isolation of arteries and isometric wire-myography

Mice were euthanized via CO_2_ inhalation and the mesentery and heart were dissected. From the left upper leg a 2 mm segment of the femoral artery was dissected. The mesentery was placed in a Petri dish fill with black silicon and ice-cold Krebs Ringer Buffer (KRB) with the following composition (in mM): 118.5 NaCl, 4.7 KCl, 2.5 CaCl_2_, 1.2 MgSO_4_, 1.2 KH_2_PO_4_, 25.0 NaHCO_3_, and 5.5 D-glucose. Second-order branches of the superior mesenteric artery (MA2) were dissected. From the heart a 1.5 - 2 mm-segment of the left descending coronary artery was dissected. Segments were mounted on a wire-myograph (Danish Myotechnology Inc, Model 620M, Aarhus, Denmark) and stretched to their optimal internal circumference as described earlier [9]. Force (mN) generated by stretch was corrected for vessel length to obtain tension values in mN/mm. Vessel length was measured in the myograph chamber with the help of a scale bar in the ocular of the stereo dissecting microscope. After an incubation period of 60 minutes, arteries were “woken up” by replacing KRB with 60 mM KCl in KRB (replacing equimolar NaCl with KCl), thus generating a stable tension after a few minutes. This tension level (minus the baseline tension) was set as 100% contraction (as % of K_60_). Cumulative concentration-response curves (CRC) were performed with phenylephrine (PHE; 0.01 – 30 μM) for MA2, and serotonin (5-HT; 0.001 – 3 μM) for coronary and femoral arteries. Cumulative CRC to acetylcholine (ACh; 0.001 – 10 μM) were assessed in contracted artery segments in the absence of any inhibitors. Endothelium-independent relaxations were assessed with CRC to the NO donor sodium nitroprusside (0.1 nM – 10 μM) in the presence of the non-selective NO synthase blocker N^ω^-nitro-L-arginine methyl ester (L-NAME; 100 μM) and the non-selective cyclo-oxygenase inhibitor indomethacin (10 μM).

### Protocol to isolate the contribution of NO in PHE-induced contractions and ACh-induced relaxing responses in MA2

The contribution of NO in PHE-induced contractions and ACh-induced relaxations was assessed by comparing the area between the curves (ABC) of the individual concentration-response curves (CRCs) in the absence and presence of the L-NAME. To rule out any contribution of vasoactive prostaglandins, all segments were treated with indomethacin in the reminder of this study.

### Protocol to isolate the contribution of EDH in ACh- and NS309-induced relaxing responses in MA2

The EDH response was blocked with selective inhibitors of small-conductance and intermediate-conductance calcium-activated potassium channels (SK_Ca_ and IK_Ca_, respectively) by UCL-1684 (1 μM) and TRAM-34 (10 μM), respectively. Both inhibitors were incubated for 30 minutes prior CRC to ACh. The EDH-mediated relaxing responses were assessed by calculating ABC between the individual CRCs to ACh and the direct endothelial K_Ca_ opener NS309 [12] in the absence and presence of TRAM-34 and UCL-1684. Another protocol was designed to assess EDH responses in MA2 that were incubated with L-NAME. The remaining relaxing responses to ACh and NS309 is an indication for the contribution of the EDH response in the absence of NO and vasoactive prostaglandins.

### Data and statistical analysis

Data are shown as mean ± SEM. Concentration-response curves were analyzed with two-way analysis of variance (ANOVA) followed by Bonferroni post-hoc test or Tukey’s test for multiple comparisons. Other values were analyzed by paired and unpaired Student’s *t* test. *P* < 0.05 was considered to be statistically significant. Sensitivity (pEC_50_) to ACh and NS309 was determined in GraphPad Prism (version 7) using nonlinear regression (variable slope with four parameters; constrains: TOP: 100, BOTTOM, 0).

## Results

### Comparable body weight after 3 months of a high fat diet regimen

The average age at euthanization was similar for all four mice groups (28 ± 1 g for lean/saline, 28 ± 1 g for obese/saline, 28 ± 1 g for lean/rhTrx, and 28 ± 2 g for obese/rhTrx). Body weight progressively increased during the 3-month high fat regimen or normal diet in both saline-infused and rhTrx-injected mice (Figure 1A). The body weight gain was roughly 15 g for mice placed on a high fat diet for 13 weeks (Figure 1B). Tail vein injection of rhTrx did not result in a statistically significantly different body weight gain compared to saline injection (Figure 1B).

**Figure 1.**
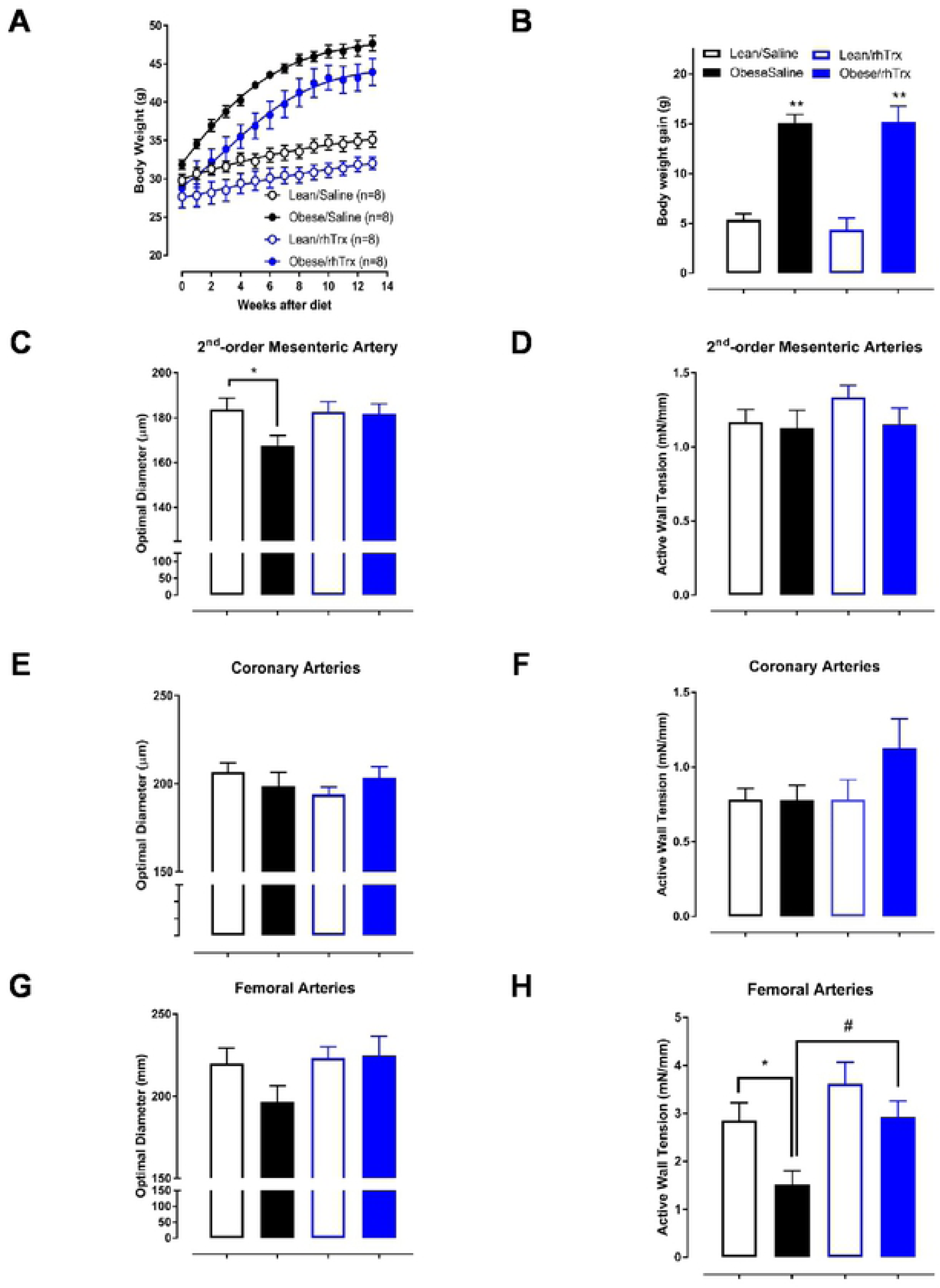
Body weight (in g) increases during the 3-month diet regimens (**A**) and body weight at the time of euthanization (**B**) for the four experimental groups. Optimal diameters (in μm) for MA2 (**C**), coronary (**E**), and femoral arteries (**G**) obtained via isometric myography. Active wall tension (in mN/mm) in response to 60 mM KCl in KRB for MA2 (**D**), coronary (**F**), and femoral arteries (**H**). Values are expressed in mean ± S.E.M. * *P* < 0.05 obese/saline versus lean/saline; # *P* < 0.05 obese/saline versus obese/rhTrx.

### Tail vein injection of rhTrx prevents high fat-induced inward remodeling of small mesenteric arteries

Second-order branches of the superior mesenteric artery (MA2) from all 4 experimental groups were mounted on an isometric wire-myograph system. Following incubation for 30 min, arteries were stretched as described in the Methods section. Optimal diameters from MA2 from obese/saline mice were statistically significantly reduced compared to their lean counterparts (Figure 1C; 168 ± 5 μm versus 184 ± 5 μm, respectively). Infusion of rhTrx prevented this inward remodeling after a high-fat diet, since optimal diameters were comparable for both lean and obese mice that were given rhTrx (Figure 1C; 183 ± 5 μm versus 182 ± 4 μm, respectively). Active wall tension (in mN/mm) in response to a depolarizing KRB solution containing 60 mM KCl were comparable in MA2 for all experimental mice groups (Figure 1D). Coronary artery optimal diameter (Figure 1E) and active wall tension (Figure 1F) were similar for the four experimental mice groups, although wall tensions tended to be larger in the obese/rhTrx group (Figure 1F). Femoral artery optimal diameter tended to be smaller in the obese/saline group compared to the other three groups (Figure 1G), but active wall tension were statistically significantly reduced compared to the other three groups (Figure 1H).

### Contractile responses are comparable for all experimental groups

Phenylephrine contracted MA2 in a concentration-dependent manner (Figure 2A). Normalized (as percentage of 60 mM KCl) concentration-response curves (CRC) showed no significant differences in both sensitivity (pEC_50_) and maximum contraction (E_max_) for PHE in all groups (Figure 1). Serotonin (5-HT) contracted coronary and femoral arteries in a concentration-dependent manner (Figure 2). Similarly to MA2, no significant differences in pEC_50_ and E_max_ were observed between the experimental groups for both artery types.

**Figure 2.**
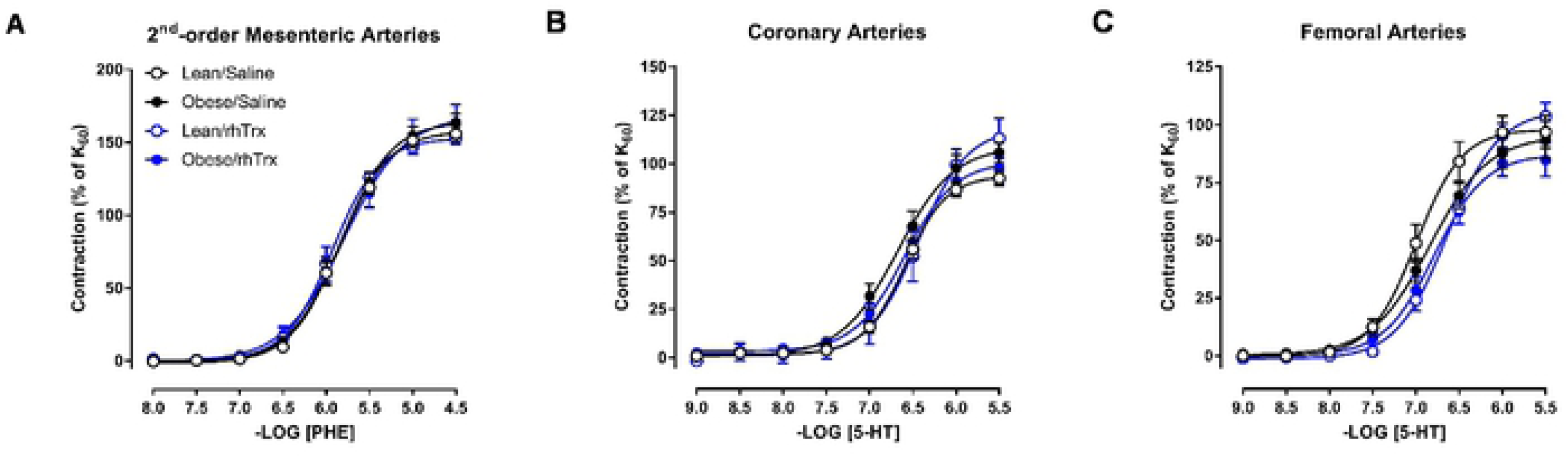
Contractile responses to the α_1_-adrenergic agonist phenylephrine (PHE; 0.01 – 30 μM) in MA2 (**A**), coronary (**B**) and femoral arteries (**C**) for the four experimental groups. Values are expressed in mean ± S.E.M.

### Acetylcholine-induced relaxing responses are impaired in MA2 and femoral arteries from obese/saline mice, but not in obese/rhTrx mice

Acetylcholine relaxed MA2, coronary and femoral arteries in a concentration-dependent manner (Figure 3A – 3C). This relaxation was diminished in MA2 and femoral arteries, but not coronary arteries, derived from obese mice injected with saline. In MA2, pEC_50_ was decreased 10-fold in obese/saline compared to lean/saline (5.45 ± 0.08 versus 6.55 ± 0.05; P < 0.05; Figure 3A). E_max_ was significantly diminished in obese/saline compared to lean/saline (58 ± 6% versus 85 ± 3%; P < 0.05; Figure 3A). ACh-induced relaxations were comparable in MA2 between lean/saline and lean/rhTrx mice (Figure 3A). Strikingly, tail vein infusion of rhTrx completely prevented the high fat-induced ACh-induced impairment, with pEC_50_ and E_max_ values similar to lean/saline values (6.25 ± 0.08 and 75 ± 7%, respectively; Figure 3A). A similar trend was observed in femoral arteries, but the differences were not as pronounced as in MA2 (Figure 3C). ACh-induced relaxations in coronary arteries were comparable for all experimental groups (Figure 3B). The observed impairments in ACh-induced relaxations were endothelium-dependent since relaxing responses to sodium nitroprusside, a NO donor with endothelium-independent relaxing properties, were not statistically significant between all groups for MA2, coronary and femoral arteries (Figure 3D – 3F).

**Figure 3.**
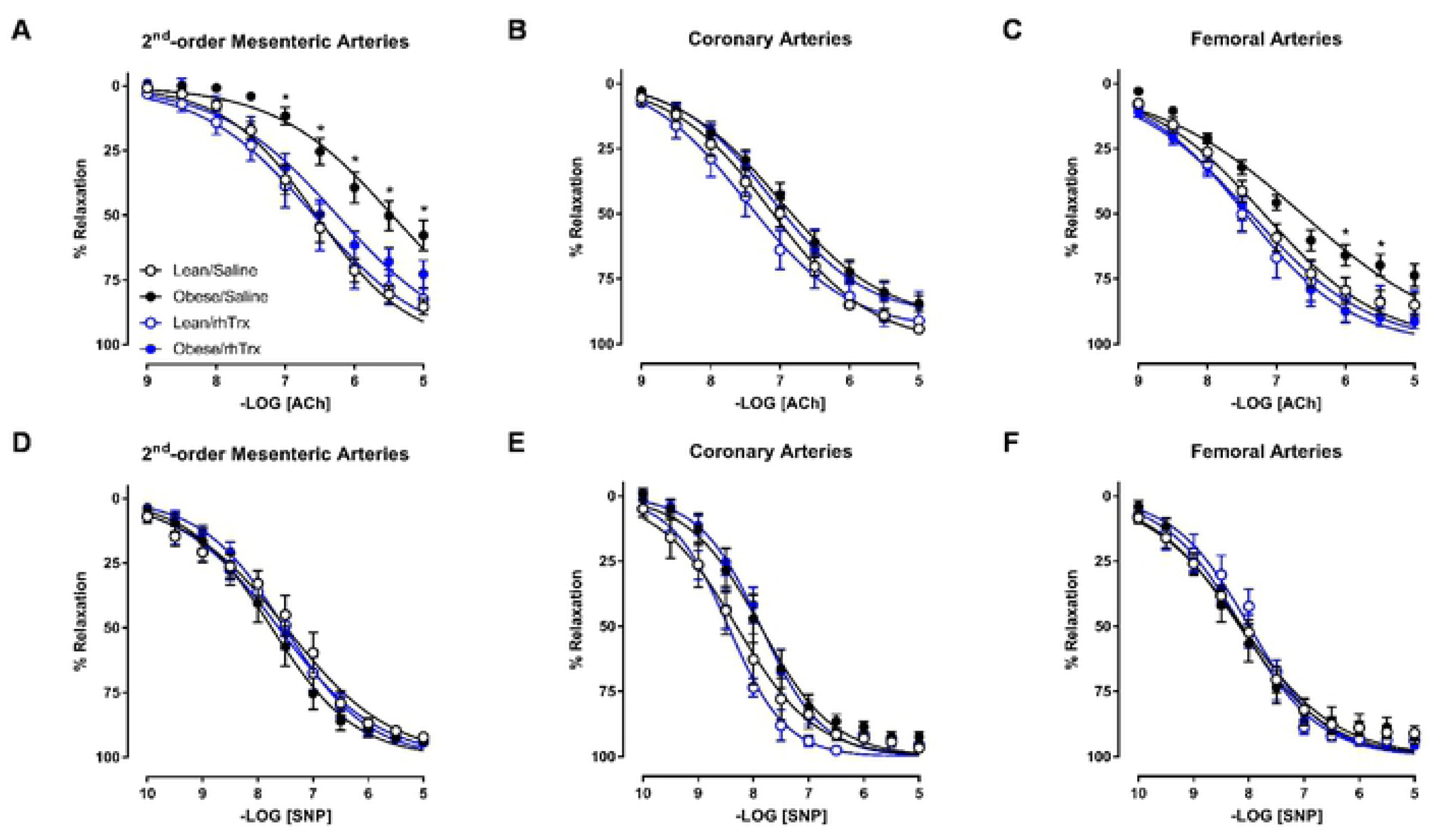
Endothelium-dependent ACh-induced relaxation in MA2 (**A**), coronary (**B**) and femoral arteries (**C**) for the four experimental groups. Endothelium-independent relaxations to the NO donor sodium nitroprusside (SNP) in MA2 (**D**), coronary (**E**) and femoral arteries (**F**) for the four experimental groups. Values are expressed in mean ± S.E.M. * *P* < 0.05 obese/saline versus the three other groups.

### NO release is diminished in MA2 from obese/saline mice, but preserved in obese/rhTrx

The suppressing role of endothelial derived NO in PHE-induced contractions was analyzed by comparing CRCs to PHE in the absence and presence of L-NAME. Figure 4A to 4D shows contractions to PHE for MA2 derived from the four groups. The area between the curves is highlighted in light blue. In a similar fashion, the role of NO in ACh-mediated relaxing responses was assessed. Figure 4E to 4H shows ACh-induced relaxing responses in the absence and presence of L-NAME with the light blue areas as area between the curves. The area under/above the curves (AUC for PHE or AAC for ACh) and the arbitrary units of the areas between the curves are depicted in Figure 4I to 4L. The surface area of the light blue areas are smaller for MA2 from obese/saline mice (Figure 4G, 4E, and 4J) compared to the other three groups, highlighting a diminished functional role of NO in vasomotor responses. More importantly, the role of NO in these responses is completely protected by tail vein injections of rhTrx (Figure 4D, 4H, and 4L).

**Figure 4.**
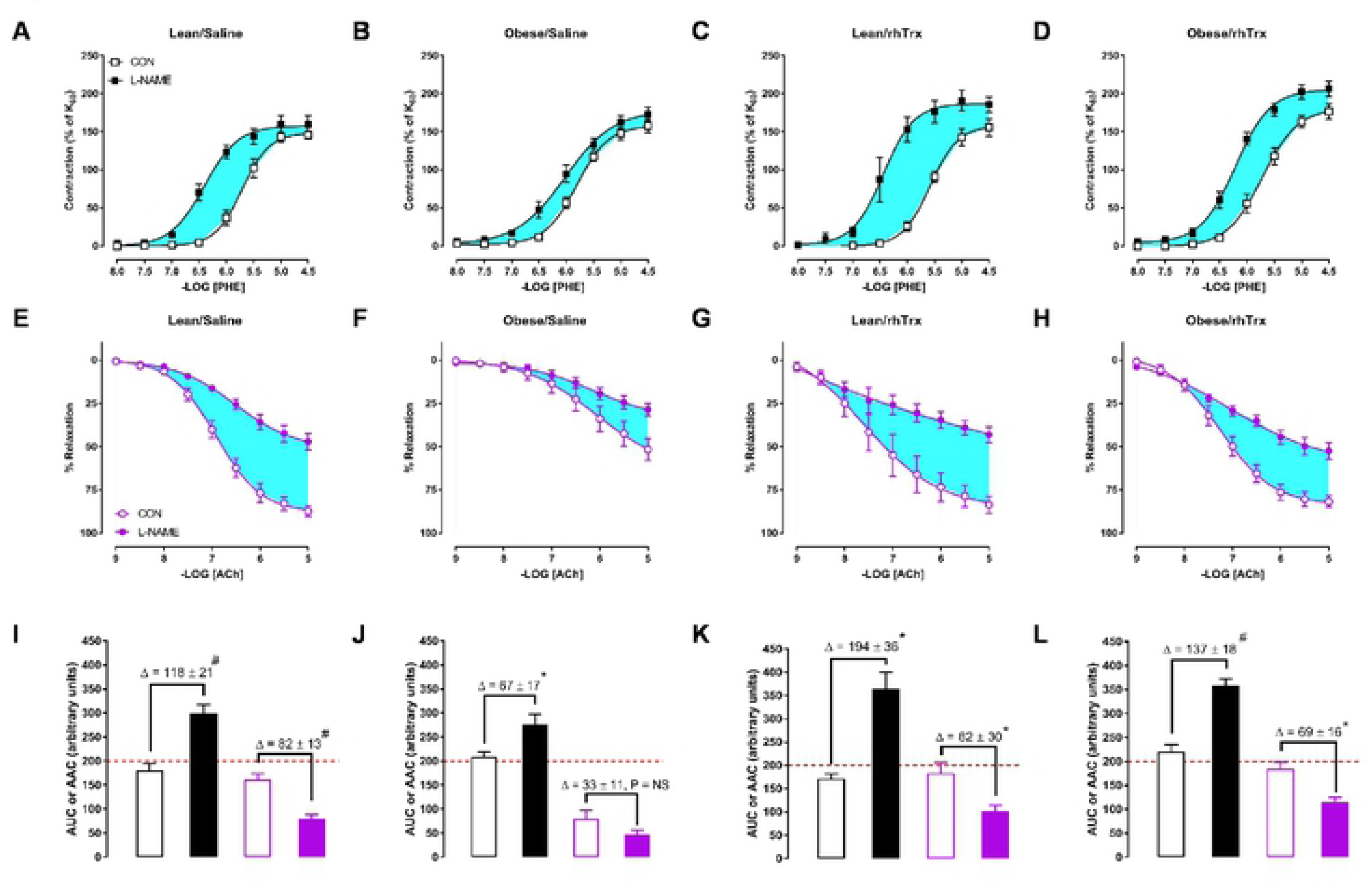
Contribution of NO in vasomotor responses. Contractile responses to the α_1_-adrenergic agonist phenylephrine (PHE; 0.01 – 30 μM) in MA2 in the absence (CON, open circles) and presence of L-NAME (100 μM, closes circles) for lean/saline (**A**), obese/saline (**B**), lean/rhTrx (**C**) and obese/rhTrx mice (**D**). Endothelium-dependent ACh-induced relaxation in MA2 in the absence (CON) and presence L-NAME (100 μM) for lean/saline (**E**), obese/saline (**F**), lean/rhTrx (**G**) and obese/rhTrx mice (**H**). Hghlighted blue areas represent the contribution of NO. Calculated area under the curve (AUC) for PHE-induced contractions (black) and calculated area above the curve (AAC) for ACh-induced (purple) responses for lean/saline (**I**), obese/saline (**J**), lean/rhTrx (**K**) and obese/rhTrx mice (**L**). Differences in AAC or AUC (Δ) are shown. All arteries were incubated with indomethacin (10 μM). Values are expressed in mean ± S.E.M. * *P* < 0.05, # *P* < 0.001, NS is not significant.

### EDH relaxing responses are diminished in obese/saline, but preserved in obese/rhTrx

The contribution of endothelial K_Ca_ channels on the EDH relaxing response was determined via two pharmacological approaches. The first approach assesses differences between CRCs to ACh and NS309 in the absence and presence of the two endothelial K_Ca_ channel blockers TRAM-34 and UCL-1684. Figure 5A to 5D shows the contribution of endothelial K_Ca_ channels that were activated indirectly via ACh-induced signaling for the four groups. The yellow highlighted areas show the ABC and are an indication of the magnitude of endothelial K_Ca_ involvement in response to ACh. The middle panel of graphs (Figure 5E to 5H) depict the contribution of these K_Ca_ channels after direct opening by NS309, with the magnitude of the EDH response highlighted in yellow. The areas under the curve (AUC) for all graphs are summarized in the bar graphs (Figure 5I to 5L). The extent of the yellow highlighted areas are shown as differences between bar heights. No significant difference between the bar graphs was observed for obese/saline mice, indicating blunted EDH response in these vessels (Figure 5J). Tail vein injections of rhTrx completely prevented the high fat-induced reduction in the EDH response (Figure 5L).

**Figure 5.**
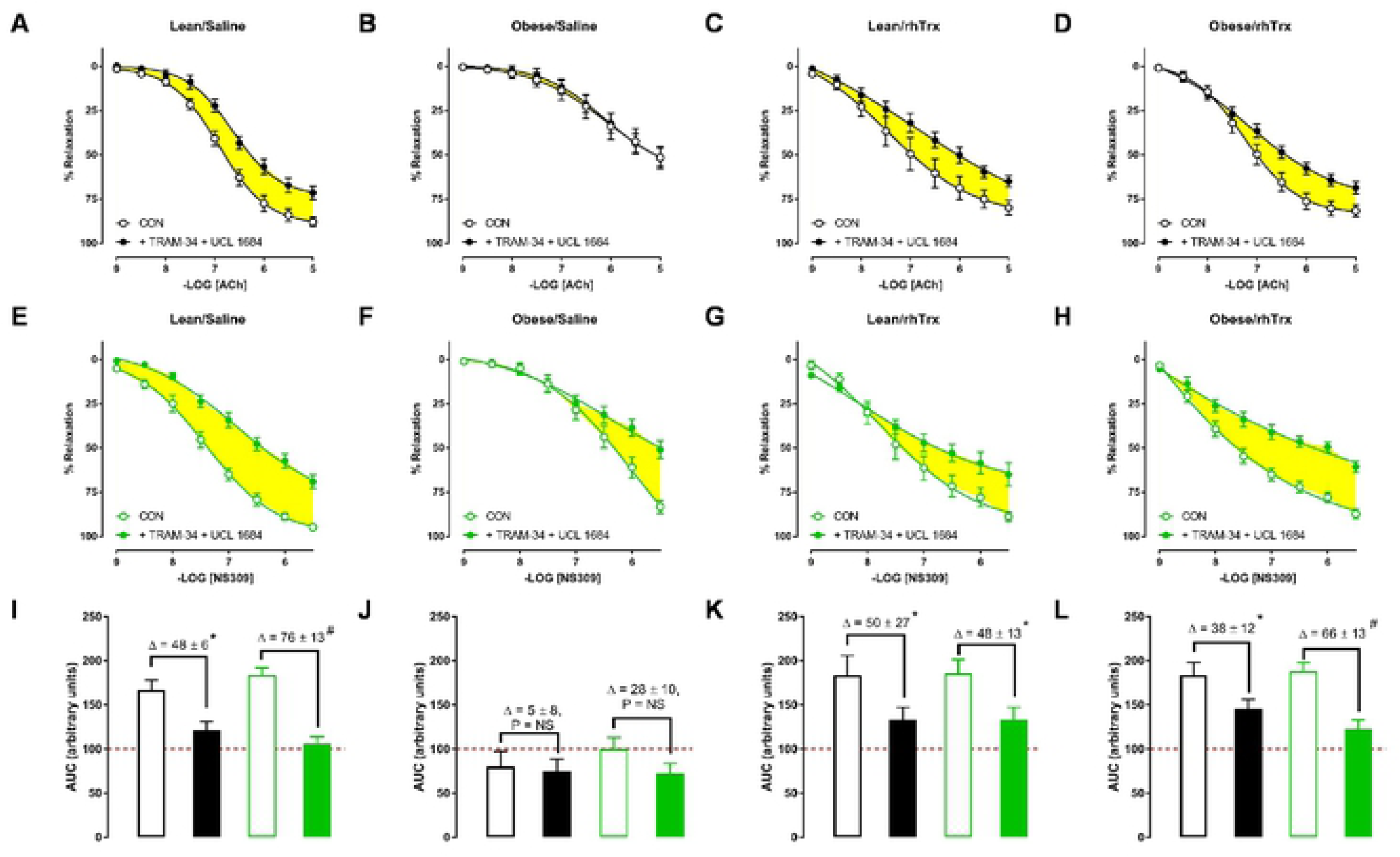
Contribution of EDH. Endothelium-dependent ACh-induced (**A** to **D**) and NS309-induced (**E** to **H**) relaxing responses in MA2 in the absence (CON, open symbols) and the presence of TRAM-34 (10 μM) + UCL-1684 (1 μM) (closed circles) for lean/saline (**A** and **E**), obese/saline (**B** and **F**), lean/rhTrx (**C** and **G**), and obese/rhTrx (**D** and **H**). Light blue areas represents the contribution of EDH via endothelial K_Ca_ channel activation. Calculated area under the curve (AUC) for ACh-induced responses (black) and NS309-induced responses (green) for lean/saline (**I**), obese/saline (**J**), lean/rhTrx (**K**) and obese/rhTrx mice (**L**). Differences in AUC (Δ) are shown. All arteries were incubated with indomethacin (10 μM). Values are expressed in mean ± S.E.M. * *P* < 0.05, # *P* < 0.001, NS is not significant.

Historically, the EDH-mediated response is analyzed in the presence of L-NAME and indomethacin. Figure 6A shows that ACh-induced EDH responses were smallest in MA2 from obese/saline mice. AAC values were significantly reduced in MA2 from obese/saline compared to the other groups (Figure 6B). Sensitivity for ACh was significantly lower in MA2 from obese/saline mice compared to the other groups (Figure 6C). NS309-induced relaxing responses were reduced in MA2 from obese/saline mice compared to the other groups (Figure 6D). AAC (Figure 6E) and sensitivity (Figure 6F) for NS309 were significantly decreased in obese/saline compared to the other groups. Again, tail vein injections of rhTrx completely prevented the high fat-induced blunted EDH response.

**Figure 6.**
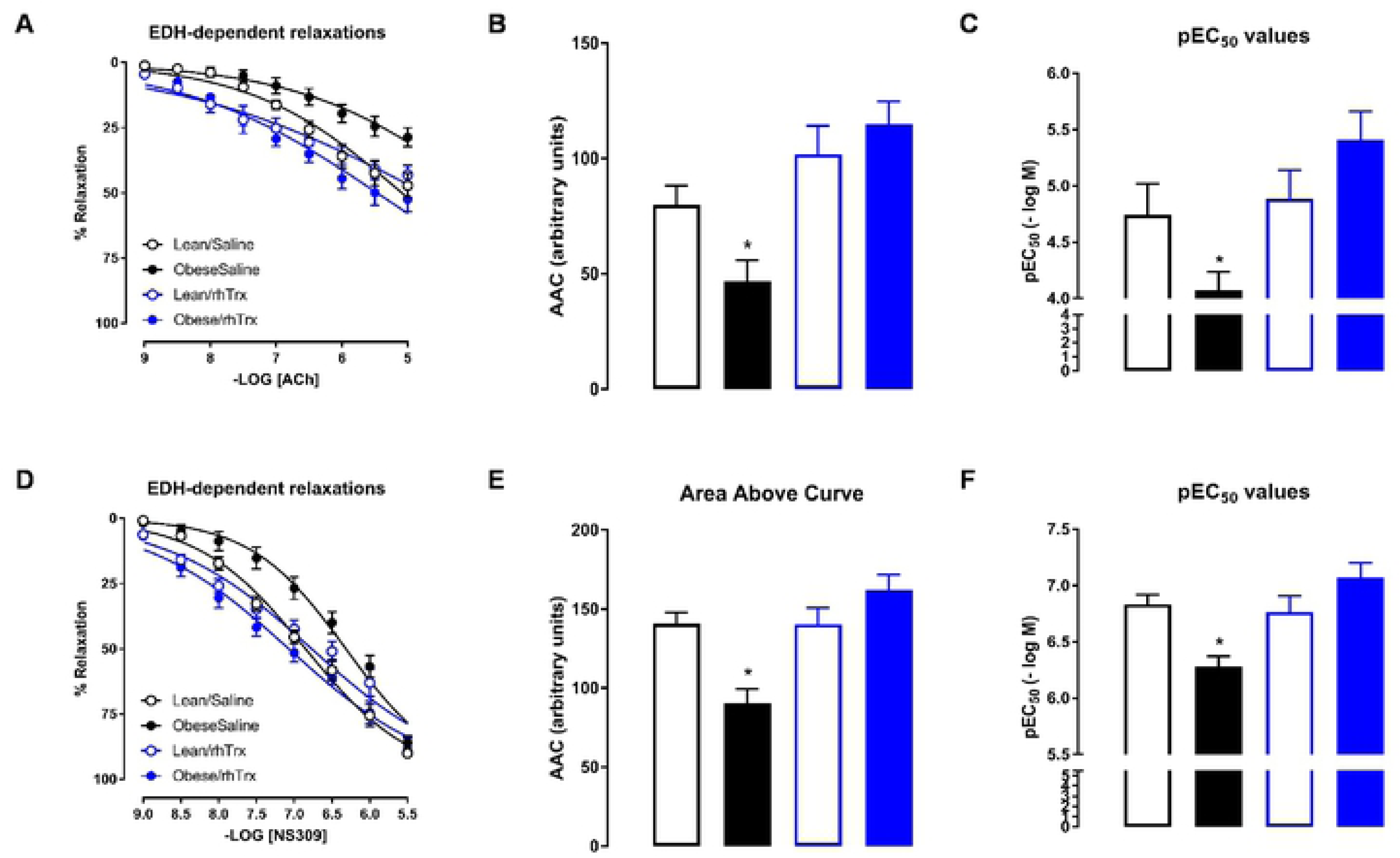
Endothelium-dependent ACh-induced (**A** to **C**) and NS309-induced (**D** to **F**) relaxing responses in MA2 in the presence of L-NAME in lean/saline (open black circles or bars), obese/saline (closed black circles or bars), lean/rhTrx (open blue circles or bars), and obese/rhTrx (closed blue circles or bars). Area above the curve are summarized in (**B** and **E**). Sensitivity (pEC_50_) are summarized in (**C** and **F**). All arteries were incubated with indomethacin (10 μM). Values are expressed in mean ± S.E.M. * *P* < 0.05 obese/saline versus all other groups.

## Discussion

The present isometric myograph functional data support the hypothesis that tail vein injections of human recombinant thioredoxin completely protects against high fat-induced endothelial dysfunction in small mesenteric and femoral arteries. In small mesenteric arteries this protection is characterized by an increased NO and EDH relaxing response, the latter via enhanced endothelial K_Ca_ channel opening.

Obesity is a major risk factor for the development of cardiovascular and metabolic complications such as hypertension and type 2 diabetes [13, 14]. This mouse strain is well suited because of its high sensitivity for obesity and type 2 diabetes in response to a high fat diet [15]. The objective of this study was not to assess plasma triglycerides and cholesterol levels, weigh subcutaneous and epididymal fat mass in order to confirm a metabolic syndrome. Here an obesity-induced diet (42% kcal from fat) was used to elicit an endothelial dysfunction in small arteries derived from C57Bl6/J mice. This was successful since small mesenteric and skeletal femoral arteries isolated from saline-injected obese mice presented classical signs of endothelial dysfunction: blunted ACh-induced relaxing responses. Judged from the comparable body weight gain during the 13 week high fat diet for saline- or rhTrx-injected obese mice, the injection of rhTrx did not have a significant impact on body mass gain. A dose of 25 mg/kg of rhTrx was chosen to inject via the tail vein. This dose was comparable to the dose used before, which resulted in detectable plasma levels and a blood pressure lowering effect in aged hypertensive mice [9].

Resistance arteries play a crucial role in blood pressure regulation and local tissue perfusion, as well as their capacity to adapt to hemodynamic changes (*e.g*. pressure, stretch and flow) [16, 17]. The optimal diameters of second-order mesenteric arteries (MA2) derived from obese/saline mice were significantly smaller than their lean counterparts, suggesting inward remodeling during obesity. Inward remodeling of small arteries has been observed in patients with essential hypertension [18], in cerebral arteries of obese rats [19], and in mesenteric arteries that underwent surgical blood flow cessation [20]. The optimal diameters were determined via horizontal stretching of isolated arteries in the wire-myograph. No pressure-myography and morphological analysis were performed to prove that the observed smaller optimal diameters were in fact the result of anatomical structural adaptations to obesity. Coronary and femoral arteries had similar optimal diameters for the four experimental groups, which suggests a regio heterogeneous effect.

Contractile responses in the absence of any inhibitors were unaltered irrespective of diet and treatment in MA2, coronary and femoral arteries. However, endothelium-dependent ACh-induced relaxing responses were impaired in MA2 and femoral arteries derived from obese/saline compared to the three counterparts. Endothelium-independent relaxing responses by the NO donor sodium nitroprusside were comparable for all groups and artery types. These observations demonstrate that the observed differences in ACh-induced responses were manifested at the level of the endothelium.

The mesenteric arterial bed is prone to high fat-induced impairment in endothelium-dependent relaxation, probably due to the close proximity of intestinal absorption of fatty acids. The small mesenteric artery is a preferred choice of resistance-sized artery for the vascular biologist due to its abundance and relative ease of dissection. Hence, endothelial dysfunction in response to obesity has been shown in murine small mesenteric arteries [21-25], but some studies did not observe an impairment in ACh-induced relaxation [26-28]. Similarly, a high-fat diet has been shown to result in either preserved endothelial function [29, 30] or impaired endothelial function in murine coronary arteries [31, 32]. Similar to this study, impaired ACh-induced relaxing responses in femoral arteries were observed in C57Bl6/J mice that received a high fat diet [30].

A pharmacological approach was used to assess the role of NO and the endothelium-dependent hyperpolarizing (EDH) relaxing response in MA2. In general, indomethacin is used in *ex vivo* vascular reactivity studies to block cyclo-oxygenases that produce vasoactive prostaglandins. Here, all MA2 were treated with indomethacin to rule out any contribution of vasoactive prostaglandins. In the present study, the main mechanism of endothelium-dependent relaxation in the murine small mesenteric artery was via NO release and EDH, which is in agreement with other studies [10, 33, 34]. In obese and saline-injected mice, both the NO- and EDH-dependent relaxation in MA2 were significantly impaired compared to lean and saline-injected mice. This observation is congruent with other studies using a high fat diet and small mesenteric arteries in the myograph [21, 22, 35]. EDH responses were assessed with both ACh (indirectly) and NS309 (directly), a non-selective endothelial IK_Ca_ and SK_Ca_ channel opener [12]. ACh increases intracellular Ca^2+^ ions in endothelial cells that activate endothelial K_Ca_ channels, whereas NS309 is an opener of these K_Ca_ channels. In murine small mesenteric arteries, the IK_Ca_ (or IK1) channel contributes mainly to ACh-stimulated Ca^2+^ dynamics [36] and genetic knockdown of IK1 reduces ACh-induced EDH response in murine mesenteric arteries [37, 38]. In agreement with these observations, the NS309-induced EDH response was inhibited by TRAM-34 and not by UCL-1684 (data not shown), confirming the importance of the IK_Ca_ channel in this species and artery type. The IK_Ca_ channel can be oxidized by hydrogen peroxide (H_2_O_2_) and other chemical cysteine thiol oxidizers, like 5,5’-dithio-bis (2-nitrobenzoic acid) (DTNB or Ellman’s reagent) or [(O-carboxyphenyl)thio]ethyl mercury sodium salt (thimerosal), to inhibit IK_Ca_ channel activity in bovine aortic endothelial cells [39]. Thiol reducing agents like dithiotreitol or reduced glutathione were able to restore the IK_Ca_ channel activity. *In vivo*, the IK_Ca_ channel can be inactivated by oxidative stress factors like obesity and hyperhomocysteinemia [40, 41]. Interestingly, the nonluminal S6 region of the IK_Ca_ channel protein, which is crucial in pore-forming, contains two adjacent cysteine residues (Cys276 and −277), which have the potential to become subject to post-translational thiol modification [42]. The above observations suggest redox modulation of the IK_Ca_ channel with an important modulatory role for thioredoxin. Using transgenic mice overexpressing human thioredoxin-1 (Trx-Tg), an enhanced EDH response was observed in MA2 compared to non-transgenic mice [10]. Furthermore, endothelial NO release was enhanced in aortae of Trx-Tg mice [9]. These observations prompted the idea of exogenously administrating rhTrx in mice in an effort to study whether rhTrx could protect against high fat-induced endothelial dysfunction.

Tail vein injection of rhTrx completely protected endothelium-dependent relaxing responses in MA2 against a high fat diet, via an increased NO release and enhanced EDH response. The latter was mediated via endothelial K_Ca_ channel activation. This observation strongly suggests that redox modulation of cysteine thiol groups on proteins regulates the release of endothelial-derived NO and EDH. Trx is also an antioxidant, because it scavenges hydroxyl radicals [43]. It could therefore be argued that the beneficial effects observed by rhTrx are attributed to its antioxidant activity. However, dominant-negative Trx mice, which express a mutant Trx without the catalytic two cysteine residues, but still possess radical scavenging properties, display blunted EDH-mediated relaxing responses in MA2 [10]. Whether Trx directly modulates K_Ca_ channel activity needs to be elucidated. A novel IK_Ca_ selective positive-gating modulator, SKA-31 [44], potentiated the EDH response in porcine coronary arteries, suggesting the important role of the IK_Ca_ channel in this artery [45]. Hence, Trx may have beneficial protective effects on coronary blood flow during conditions that reduce expression of endothelial K_Ca_ channels, such as diabetes [46-48] and hypertension [49-51].

In conclusion, tail vein injection of rhTrx completely protected obese mice from high fat-induced endothelial dysfunction. In addition, compared to lean/saline-injected mice, NO and EDH-mediated responses were enhanced in MA2 from obese/rhTrx-injected mice. These vasoprotective actions of Trx may provide a promising therapeutic potential in the combat of oxidative stress-linked pathologies.

## Acknowledgement

Funding for this study was supported by a New Investigator Award (2019) from the American Association of Colleges of Pharmacy.

